# Evaluation of mRNA-LNP and adjuvanted protein SARS-CoV-2 vaccines in a maternal antibody mouse model

**DOI:** 10.1101/2023.04.12.536590

**Authors:** Ross N. England, Elizabeth M. Drapeau, Reihaneh Hosseinzadeh, Drew Weissman, Scott E. Hensley

## Abstract

Maternal antibodies (matAbs) protect against a myriad of pathogens early in life; however, these antibodies can also inhibit *de novo* immune responses against some vaccine platforms. Severe acute respiratory syndrome coronavirus 2 (SARS-CoV-2) matAbs are efficiently transferred during pregnancy and protect infants against subsequent SARS-CoV-2 infections. It is unknown if matAbs inhibit immune responses elicited by different types of SARS-CoV-2 vaccines. Here, we established a mouse model to determine if SARS-CoV-2 spike-specific matAbs inhibit immune responses elicited by recombinant protein and nucleoside-modified mRNA-lipid nanoparticle (mRNA-LNP) vaccines. We found that SARS-CoV-2 mRNA-LNP vaccines elicited robust *de novo* antibody responses in mouse pups in the presence of matAbs. Recombinant protein vaccines were also able to circumvent the inhibitory effects of matAbs when adjuvants were co-administered. While additional studies need to be completed in humans, our studies raise the possibility that mRNA-LNP-based and adjuvanted protein-based SARS-CoV-2 vaccines have the potential to be effective when delivered very early in life.

## Introduction

Since late 2019, severe acute respiratory syndrome coronavirus 2 (SARS-CoV-2) has caused a global pandemic that has prompted the rapid development of multiple vaccines, including vaccines that employ the mRNA-lipid nanoparticle (mRNA-LNP)-based platform. These vaccines have been shown to be efficacious and safe in adults^1,2^ and children^3-5^ and have been authorized for use in persons 6 months and older in the United States. Because no SARS-CoV-2 vaccines are authorized for use in infants less than 6 months of age, this age group relies on aggregate population immunity and circulating maternal antibodies (matAbs) for protection from SARS-CoV-2 infection.

MatAbs provide early life protection against a variety of pathogens. However, matAbs also can interfere with seroconversion in response to infections and vaccines,^6,7^ including live-attenuated vaccines (e.g., measles, mumps), acellular and whole cell bacterial (e.g., pertussis), inactivated viral vaccines (e.g., IPV, influenza virus), and recombinant protein subunit vaccines (e.g., hepatitis B virus). MatAb interference of vaccine-elicited responses can negatively impact efficacy of early life vaccination programs and potentially expose infants to a window of opportunity for infection while awaiting effective vaccination and consequent *de novo* IgG responses. Our mechanistic understanding of matAb interference remains incomplete; available evidence suggests that matAbs can mask antigen and inhibit infant naïve B cell activation via FCγRIIB binding.^6,8-10^

In humans, SARS-CoV-2 matAbs are efficiently transferred in utero^11,12^ and remain detectable at 6 months of age in over 50 percent of infants born to mothers vaccinated during pregnancy^13^. This suggests that any efforts at vaccinating infants younger than 6 months of age might be limited by matAb interference. It is unknown if matAbs inhibit *de novo* immune responses elicited by SARS-CoV-2 mRNA-LNP vaccines. Our group previously demonstrated that mRNA-LNP-based influenza vaccines partially overcome matAb interference in mice,^14^ which raises the possibility that SARS-CoV-2 mRNA-LNP vaccines may remain effective in the presence of SARS-CoV-2-specific matAbs. Here, we established a SARS-CoV-2 matAb mouse model to evaluate if SARS-CoV-2 mRNA-LNP and recombinant protein vaccines are inhibited by matAbs.

## Results

### mRNA-LNP vaccines and adjuvanted recombinant protein vaccines elicit SARS-CoV-2 IgG responses in mouse pups

We first evaluated antibody responses to multiple SARS-CoV-2 vaccines in weanling mice in the absence of maternal antibodies. We vaccinated matAb-negative C57BL/6 mouse pups at weaning (∼21 days of life) with SARS-CoV-2 mRNA-LNP (1μg, 5μg, or 10μg dose), recombinant spike protein (RP) diluted in PBS (RP-PBS; 5μg dose), RP adjuvanted with empty LNP (RP-LNP, 5μg dose), or RP adjuvanted with the M59-like adjuvant Addavax™ (RP-Ad; 5μg dose). We obtained serum samples and measured SARS-CoV-2 full-length spike protein-specific IgM and IgG by ELISA over time (Figure 1a). Vaccine immunogens in our study consisted of full-length spike protein from the ancestral Wuhan Hu-1 strain of SARS-CoV-2, with diproline substitution to maintain pre-fusion conformation.

**Figure 1.**
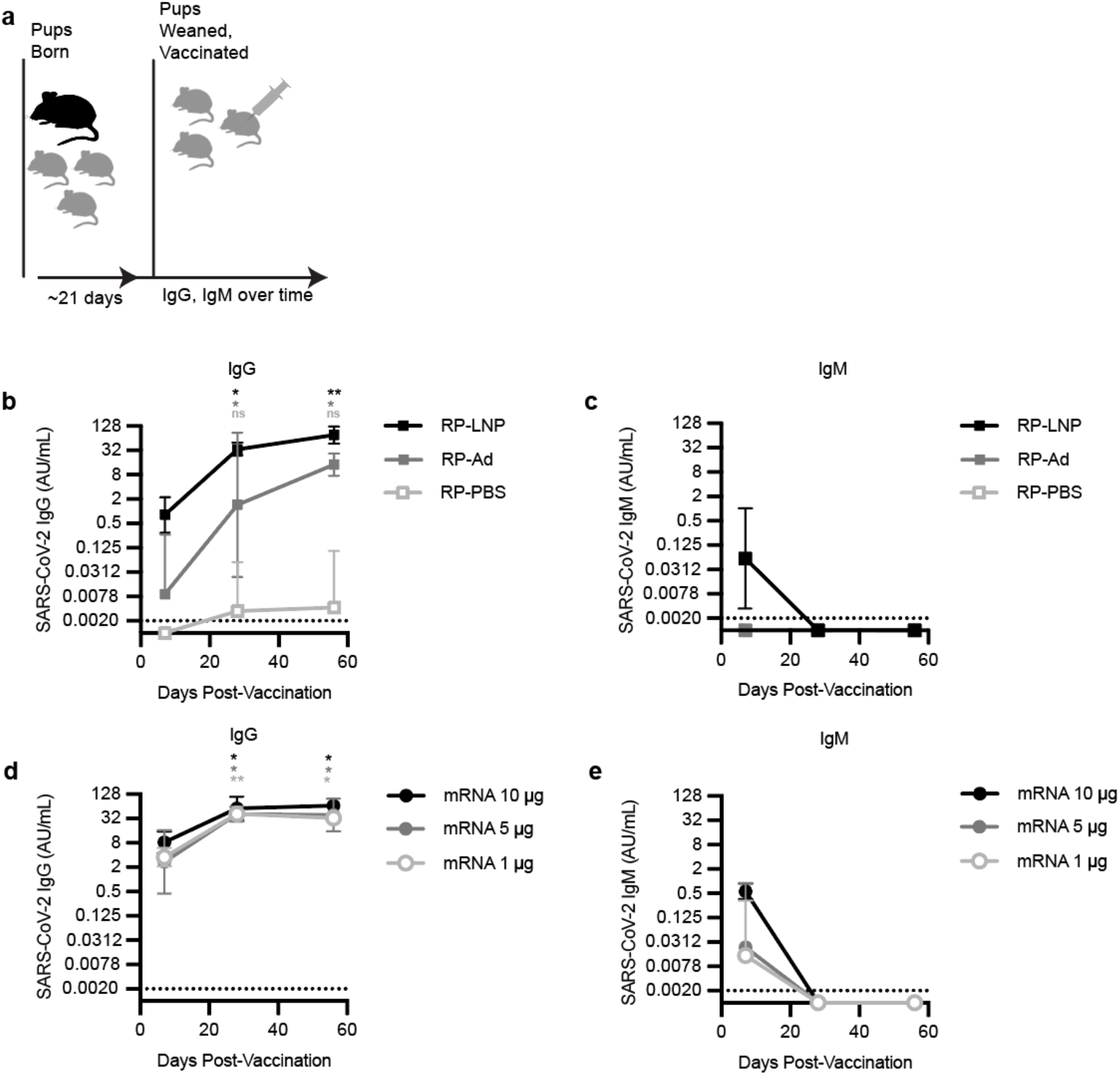
mRNA-LNP vaccines and adjuvanted protein subunit vaccines elicit SARS-CoV-2 IgG responses in mouse pups. (A) The experimental design is shown; Mouse pups were inoculated at weaning with 5μg SARS-CoV-2 recombinant protein (RP) vaccine (B, C) adjuvanted with PBS (RP-PBS), AddavaxTM (RP-Ad), or LNP (RP-LNP) or SARS-CoV-2 mRNA-LNP vaccine (D, E) at 1μg, 5μg, or 10 μg. (B-E) Sera were collected from mouse pups at indicated time points after weaning/vaccination and SARS-CoV-2 full length Spike (S) protein-specific IgG concentrations (B, D) and IgM concentrations (C, E) were measured by ELISA. Each point represents the geometric mean with error bars indicating 95% confidence interval (CI) of the geometric mean. Mouse groups were n = 5 (RP vaccinated groups) or n = 6 (mRNA vaccinated groups). Sample IgG and IgM concentrations were plotted to a standard curve of known concentrations of a S-specific IgG monoclonal antibody and are reported as arbitrary units (AU)/mL. Data are shown as geometric mean concentrations with 95% confidence intervals. All panels show results of one experiment that is representative of two independent biological replicates. Group geometric mean IgG concentrations were compared at 4 weeks and 8 weeks post-vaccination to corresponding within-group 1 week concentrations by repeated measures (RM) one-way ANOVA with Tukey’s post hoc test (* indicates p< 0.05, ** indicates p<0.01).

The RP vaccine was not immunogenic in mice without co-administering adjuvant (Figure 1b-c). In mice receiving RP-PBS, spike-specific IgG was not detected in any mice at 1 week post-vaccination and was detected in only a single mouse at 4 and 8 weeks post-vaccination (Figure 1b). Both adjuvanted RP vaccines (RP-Ad and RP-LNP) as well as mRNA-LNP vaccines (at all three doses) elicited spike-specific IgG responses in weanling mice (Figures 1b and 1d, respectively). In mice receiving RP-Ad, spike-specific IgG was detected in most mice at 1 week post-vaccination and increased significantly compared to this 1 week baseline in all mice at 4 and 8 weeks post-vaccination (Figure 1b). In mice receiving RP-LNP, spike-specific IgG was detected in most mice at 1 week post-vaccination and increased significantly in all mice at 4 and 8 weeks post-vaccination (Figure 1b). In groups receiving mRNA-LNP, spike-specific IgG was detectable at 1 week and increased significantly compared to each respective baseline at 4 and 8 weeks post vaccination (Figure 1d).

In mice receiving RP-PBS or RP-Ad, spike-specific IgM was not detected at 1, 4, or 8 weeks post-vaccination (Figure 1c). In most mice receiving RP-LNP, spike-specific IgM was detectable at 1 week post-vaccination before waning below the threshold of detection at 4 and 8 weeks post-vaccination (Figure 1c). Spike-specific IgM was detected at 1 week post-vaccination in most mice receiving 1μg and 5μg mRNA-LNP and was detectable in all mice receiving 10μg mRNA-LNP before waning to undetectable levels at 4 and 8 weeks post-vaccination (Figure 1e).

These experiments show that all SARS-CoV-2 mRNA-LNP vaccines and adjuvanted recombinant protein vaccines elicit antibody responses in weanling mouse pups. For ongoing experiments, 1μg mRNA-LNP was used for vaccination of weanling pups as it was the lowest dose of mRNA-LNP eliciting comparable spike-specific IgM and IgG responses compared to adjuvanted RP vaccines.

### Establishment of a SARS-CoV-2 matAbs mouse model

We next established a mouse model of SARS-CoV-2 matAb transfer. We determined the transfer efficiency and longevity of SARS-CoV-2 spike-specific matAbs following maternal vaccination. For these experiments, we vaccinated adult pregnant female C57BL/6 mice with SAR-CoV-2 spike mRNA-LNP at varying doses (0.1-5μg) 5 days after introducing unvaccinated male breeding partners. These mice were then allowed to deliver pups. Because matAbs are transferred to mouse offspring both in utero and via milk,^6^ we collected serum from pups at weaning (∼ 21 days of life) and at intervals over 9 weeks post-weaning and measured SARS-CoV-2 full-length spike protein-specific IgG by enzyme-linked immunosorbent assay (Figure 2a. Vaccinated dams transferred SARS-CoV-2 spike-specific matAbs to pups in a dose-dependent manner. All pups born to vaccinated dams had detectable spike-specific IgG and these spike-specific matAbs waned to undetectable levels over time in the 0.1μg, 1μg, and 5μg vaccine dose groups (Figure 2b, 0.1μg AUC 25.00 AU, 95% CI 23.03 - 26.97 AU; 1 μg AUC 156.4 AU, 95% CI 124.1 - 188.8 AU; 5 μg AUC 382.6 AU, 95% CI 306.3 - 458.8 AU). MatAb half-life was similar in all groups (Figure 2b). On day of weaning, dams had higher spike-specific IgG levels than pups (Figure 2c). Offspring:dam pairs had placental IgG transfer ratios of ∼ 0.57 (mean 0.57, range 0.20 to 1.53). These experiments demonstrate that SARS-CoV-2 spike-specific maternal antibodies are transferred to pups and wane over time.

**Figure 2.**
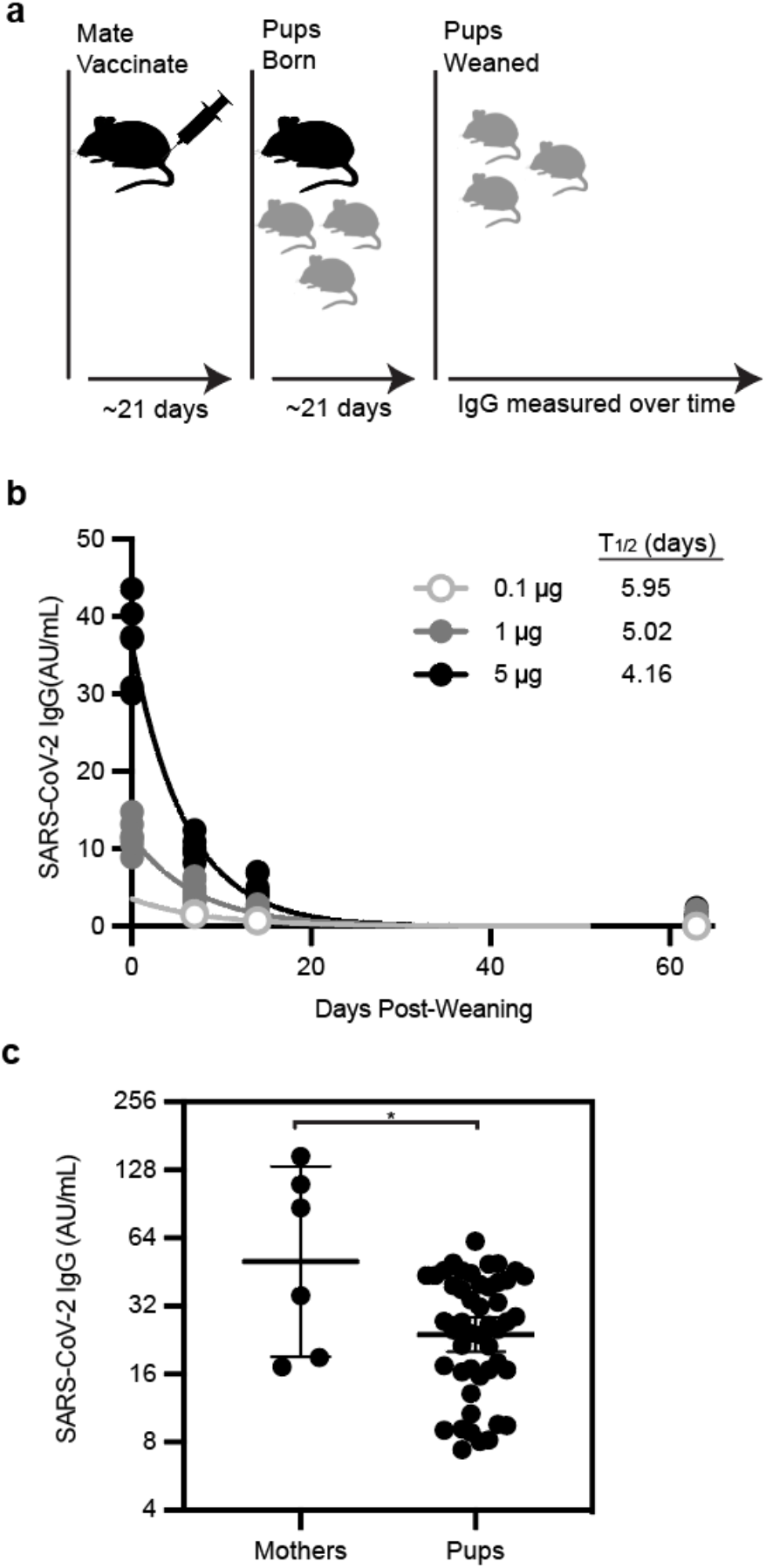
MatAbs against SARS-CoV-2 are transferred to mouse pups and wane over time. (A) The experimental design is shown; Female mice were mated with males and then were inoculated at day 5 after introduction of males with SARS-CoV-2 mRNA-LNP vaccine at 0.1μg, 1μg, or 5μg. (B-C) (B) Sera were collected from mouse pups at indicated time points after weaning/vaccination and SARS-CoV-2 full length Spike (S) protein-specific IgG concentrations were measured by ELISA. One phase-decay was fitted to IgG data (R^2^ > 0.90 for all groups) with each point representing a single mouse pup and each line representing the decay curve for one dose group (0.1μg n=2; 1μg n=12; 5μg n=7). (C) Dams were vaccinated with 5μg mRNA-LNP and serum was collected from dams (n = 6) and pups (n = 48) at day of weaning. Each point represents the geometric mean with error bars indicating 95% confidence interval (CI) of the geometric mean. Sample IgG concentrations were plotted to a standard curve of known concentrations of a S-specific IgG monoclonal antibody and are reported as arbitrary units (AU)/mL. Data are shown as geometric mean concentrations with 95% confidence intervals (* indicates p< 0.05).

### SARS-CoV-2 mRNA-LNP vaccine and adjuvanted protein subunit vaccine IgG responses are not inhibited by SARS-CoV-2-specific matAbs

We tested different SARS-CoV-2 vaccines in our newly established SARS-CoV-2 matAb mouse model. We vaccinated matAb-positive and matAb-negative C57BL/6 mouse pups at weaning (∼21 days of life) with SARS-CoV-2 mRNA-LNP (1μg dose), RP adjuvanted with LNP (RP-LNP; 5μg dose), RP adjuvanted with the MF59-like adjuvant Addavax™ (RP-Ad; 5μg dose), or PBS and measured SARS-CoV-2 full-length spike-specific IgG by ELISA over time (Figure 3a). We did not test RP without adjuvant in these experiments since we previously found that the unadjuvanted RP vaccine was poorly immunogenic even in the absence of matAbs (Figure 1b-c).

**Figure 3.**
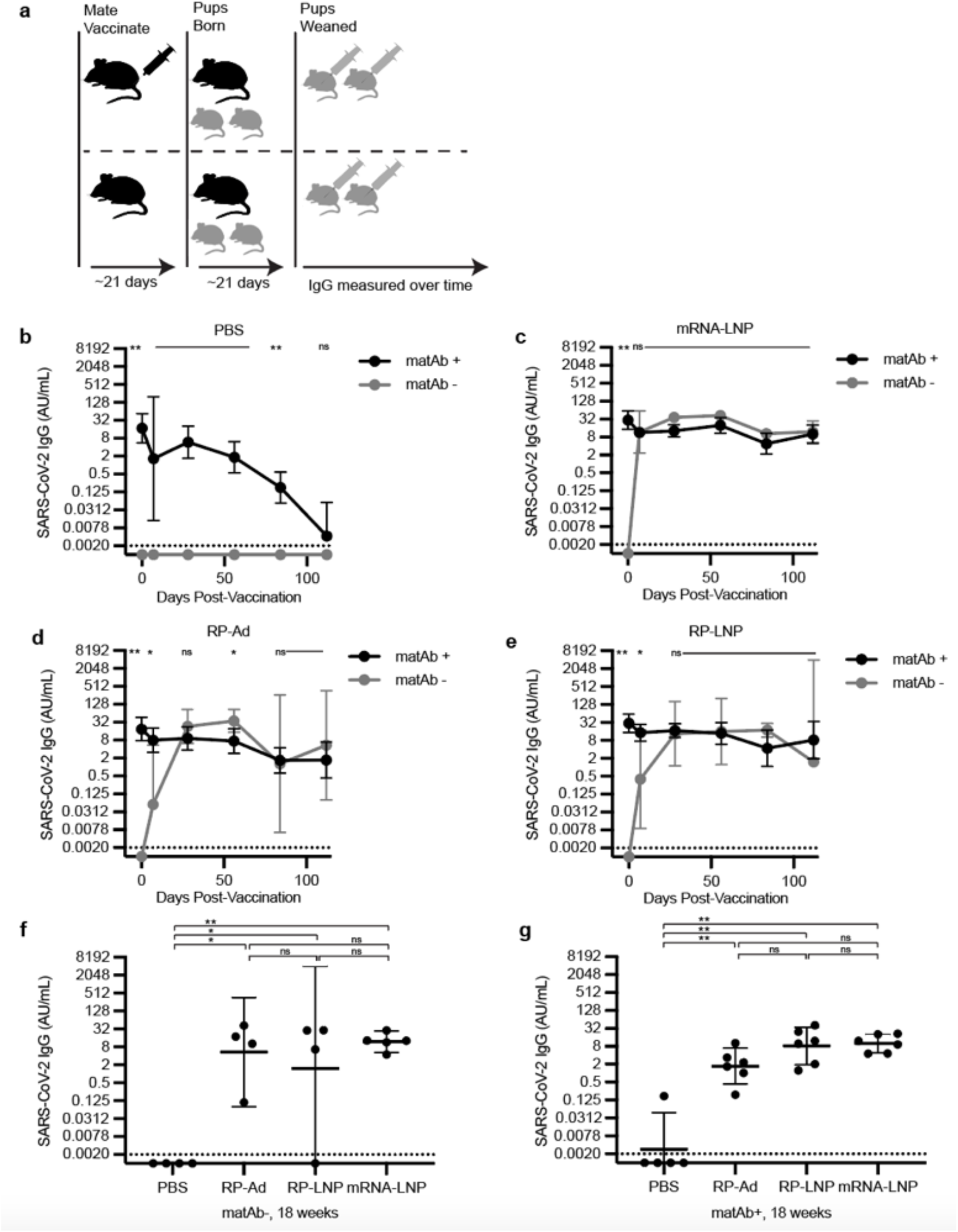
SARS-CoV-2 mRNA-LNP vaccines and adjuvanted protein subunit vaccine-elicited IgG responses are not inhibited by SARS-CoV-2-specific matAbs. (A) The experimental design is shown; Female mice were mated with males and then were inoculated at day 5 after introduction of males with SARS-CoV-2 mRNA-LNP vaccine (5μg) or PBS and pups were vaccinated at weaning with PBS, SARS-CoV-2 recombinant protein (RP) with Addavax (RP-Ad) or empty lipid nanoparticle (RP-LNP), or SARS-CoV-2 mRNA vaccine (mRNA-LNP). (B-E) Sera were collected from mouse pups at indicated time points after weaning/vaccination and SARS-CoV-2 full length Spike (S) protein-specific IgG concentrations were measured by ELISA. For each vaccine condition (B, PBS; C, mRNA-LNP; D, RP-Ad; E, RP-LNP; n = 4-6 mice per group) at each time point, geometric mean IgG concentrations were compared between the matAb+ and matAb-groups by one-way ANOVA with post hoc Šidák test. Each point represents the geometric mean with 95% confidence intervals. (F-G) Column graphs depict SARS-CoV-2 IgG concentrations for matAb-(F) and matAb+ (G) mice at 18 weeks post-vaccination for each vaccine condition, with each point representing a single mouse and central bars representing geometric mean concentration and 95% CI. Group geometric mean IgG concentrations were compared by one-way ANOVA with Tukey’s post hoc test with significance level of group comparison indicated by nested brackets. For all panels, sample IgG concentrations were plotted to a standard curve of known concentrations of a SARS-CoV-2 IgG monoclonal antibody and are reported as arbitrary units (AU)/mL with samples for which IgG was below the threshold of detection (TOD, represented by horizontal dotted lines) imputed to half the TOD (represented by the x axis). Central points/bars and error bars for all panels represent geometric mean concentrations with 95% confidence intervals. Significance levels for statistical hypothesis testing are indicated (* indicates p< 0.05, ** indicates p<0.01, ns indicates p≥0.05). All panels show results of one experiment that is representative of two independent biological replicates.

In the matAb-negative (matAb-) group vaccinated with PBS, spike-specific IgG was not detected at any time point. In the matAb-positive (matAb+) group vaccinated with PBS, spike-specific IgG waned to near the threshold of detection at 18 weeks (126 days) post-weaning (Figure 3b), consistent with the matAb decay rates that we observed in earlier experiments (Figure 2). In matAb+ mice prior to vaccination, spike-specific IgG was detected on day 0 (at weaning) in all mice and was significantly elevated compared to within-group matAb-mice, none of which had detectable IgG on day 0 (Figures 3c-e). At 7 days post-vaccination, matAb-mice receiving SARS-CoV-2 mRNA-LNP vaccines developed detectable IgG antibodies at concentrations similar to matAb+ mice (Figure 3c) and these remained detectable at similar levels until the last point of sampling at 126 days post-vaccination (Figure 3c). Mice vaccinated with adjuvanted recombinant protein SARS-CoV-2 vaccines exhibited similar trends albeit with antibody concentrations rising to similar levels between matAb+ and matAb-groups occurring at 28 days post-vaccination. At 28 days post-vaccination, matAb-mice receiving either SARS-CoV-2 RP-LNP or SARS-CoV-2 RP-Ad vaccines developed detectable IgG antibodies at concentrations similar to matAb+ mice (Figure 3e,d) and these remained detectable at similar levels until the last point of sampling at 126 days post-vacation (Figure 3e,d).

At 18 weeks (126 days) post-vaccination, spike-specific IgG concentrations in all vaccinated mouse groups were significantly higher than PBS control in both matAb+ and matAb-arms (Figures 3f and 3g, respectively). Comparing between matAb-groups, spike-specific IgG concentrations were not significantly different between mRNA-LNP, RP-Ad, and RP-LNP. Similarly, in matAb+ groups, spike-specific IgG concentrations were not significantly different between mRNA-LNP, RP-Ad, and RP-LNP. These experiments demonstrate that mRNA-LNP and adjuvanted recombinant SARS-CoV-2 vaccines elicit robust IgG responses in weanling mouse pups in the presence of SARS-CoV-2 spike-specific matAbs at the time of vaccination.

## Discussion

MatAb interference has long been an obstacle to effective immunization of newborns and young infants using conventional vaccines.^6,7^ Thus far, data suggest that severe SARS-CoV-2 infections are less common in newborns and young infants, with lower morbidity and mortality than other common pediatric respiratory viruses (e.g., influenza, respiratory syncytial virus),^20^ which has lessened the urgency to test SARS-CoV-2 vaccines in infants less than 6 months of age. However, rapid emergence of SARS-CoV-2 variants of concern and the ever-present possibility of a newly emerging pandemic coronavirus, either of which could potentially have higher impact in infants, raise the question of whether this age group can be successfully vaccinated against SARS-CoV-2 in the presence of matAbs. In this study, we established a mouse model of SARS-CoV-2-specific matAb transfer and found that both SARS-CoV-2 mRNA-LNP vaccines and adjuvanted RP spike vaccines are effective at eliciting *de novo* antibody responses in the presence of spike-specific matAbs.

The American College of Obstetrics and Gynecology (ACOG) and the Society for Maternal-Fetal Medicine (SMFM) both recommend that incompletely vaccinated pregnant women receive intragestational vaccination with an approved SARS-CoV-2 vaccine.^21,22^ Given these recommendations, we developed a mouse model involving SARS-CoV-2 matAb transfer following intragestational vaccination. We found that vaccination of pregnant dams resulted in a dose-responsive transfer of matAbs to pups that waned over time with a half-life of ∼4-6 days. Efficiency of matAb transfer to pups, measured as dam:offspring IgG ratio, was ∼ 0.57, which is similar to placental transfer ratios of SARS-CoV-2 matAbs in human mother-infant pairs in which the mother received a single dose of SARS-CoV-2 mRNA vaccine < 30 days prior to delivery.^23^ Notably in humans, SARS-CoV-2 matAb levels have been shown to be higher after maternal vaccination compared to after maternal infection but maternal vaccination leads to slightly lower placental transfer ratios.^23^ Further studies should evaluate if matAbs elicited by SARS-CoV-2 infections versus vaccinations have different inhibitory capacities in mouse models.

Nucleoside-modified mRNA-LNP vaccine technology is a safe, versatile, scalable platform that has the potential to overcome challenges of conventional vaccine technologies, and prior work from our laboratory showed that mRNA-LNP influenza vaccines (encoding influenza virus hemagglutinin) were able to partially overcome matAb interference observed with conventional split inactivated influenza vaccines.^14^ Extrapolating from these findings, we hypothesized that SARS-CoV-2 mRNA vaccines would remain effective in mice in the presence of SARS-CoV-2-specific matAbs. Consistent with this, we observed that SARS-CoV-2 mRNA-LNP vaccines evoked *de novo* IgG responses that were similar between mice with and without SARS-CoV-2 spike-specific matAbs. Since our previous experiments demonstrated that matAbs inhibit *de novo* responses elicited by adjuvanted influenza vaccines in mice,^14^ we were surprised to observe that adjuvanted SARS-CoV-2 RP vaccines were effective at eliciting *de novo* IgG in the presence of SARS-CoV-2-specific matAbs. While unlikely, it is possible that intrinsic properties of the SARS-CoV-2 spike protein (rather than a property of the specific vaccine platform and adjuvants) allows it to be immunogenic in the presence of matAbs. Further studies should explore how different adjuvants and vaccine platforms may be able to overcome the inhibitory effects of matAbs using a variety of different immunogens.

It remains mechanistically unclear how mRNA-LNP vaccines overcome the inhibitory effects of matAbs. Our previous studies with influenza vaccines suggest that long-lived germinal centers induced by mRNA-LNP vaccines likely contribute to overcoming matAb interference.^14^ Recent data suggest the empty LNP has potent adjuvant effects when combined with protein subunit vaccines and these adjuvant effects likely substantially contribute to the impressive immunogenicity of the mRNA-LNP vaccine platform.^24^ Our results in the current study suggest that novel lipid-based adjuvants, including empty LNP and MF-59-like adjuvants, may increase the effectiveness of protein subunit vaccines in the presence of maternal antibodies. This raises the possibility that the adjuvant properties of LNP contribute to mRNA-LNP vaccine effectiveness in the presence of matAbs.

There is not yet an approved SARS-CoV-2 vaccine for infants under 6 months of age and thus the only current means of immunity for this age group is passive protection from transferred matAbs, a process that has historically limited active immunization of young infants via matAb interference with vaccine antigen. Our results suggest that SARS-CoV-2 mRNA-LNP vaccines currently in use are likely to be effective in infants despite the presence of SARS-CoV-2-specific matAbs and add to a body of literature suggesting that nucleoside-modified mRNA-LNP vaccines may be an overall solution to problems associated with matAb interference.

## Methods

### Study design

The study objectives were to determine the ability of SARS-CoV-2 mRNA-LNP and protein subunit vaccines to elicit *de novo* antibody responses in the presence of SARS-CoV-2 spike-specific matAbs. Mouse pups with and without SARS-CoV-2 spike-specific matAbs were vaccinated with SARS-CoV-2 spike mRNA-LNP or adjuvanted recombinant protein subunit vaccine and antibody responses were quantitated over time. The number of pups in each experiment varied due to differences in litter size. Both male and female pups were used in all experiments. No mice were excluded from experiments and outliers were included in all analyses. Investigators were not blinded. Two biological replicates were performed for each experiment unless otherwise noted.

### Mouse model

C57BL/6 mice were purchased from Charles River Laboratories (Wilmington, MA) and bred in-house. Female mice (ages 6-8 weeks) were mated in trio with males of the same strain. On day five after introduction of males, males were removed and female mice received intramuscular (i.m.) vaccination with SARS-CoV-2 spike mRNA-LNP (0.1μg, 1μg, 5μg, or 10μg in 50 μL PBS) or vehicle. Pregnant females were singly housed and allowed to have pups. Pups were weaned at ∼21 days of age (range, 19 – 22 days) and vaccinated at the time of weaning. All mouse experiments were approved by the Institutional Animal Care and Use Committees of the Wistar Institute and the University of Pennsylvania. Sample size for each experiment was determined based on similar previous experiments.

### Serum collection

Blood was collected at indicated time points by submandibular puncture into 1.1 mL Z-Gel tubes (Sarstedt, Numbrecht, Germany) using 5-5.5mm lancets (MEDIpoint, Mineola, NY). Sera were isolated and stored at -80°C prior to thawing for analysis and thereafter stored at 4°C.

### Vaccinations

mRNA-LNP vaccines were diluted at indicated RNA amounts (0.1 – 10 μg) in PBS. Recombinant spike protein subunit vaccines were prepared by diluting 5 μg recombinant spike protein in 50 μL diluent; diluent for Addavax™-adjuvanted vaccines consisting of 25 μL PBS and 25 μL Addavax™ (Invivogen, San Diego, CA) and diluent for lipid nanoparticle-adjuvanted vaccines consisted of a concentration of LNP corresponding to the mRNA-LNP vaccine brought to a total volume of 50 μL with PBS. All vaccines were stored on ice after dilution and prior to administration. Vaccines (50 μL) were administered by intramuscular (i.m.) injection into the upper hind leg. Vaccinations were performed under isoflurane anesthesia (4 – 5% in oxygen).

### mRNA-LNP production

Nucleoside-modified (with m1Ψ) mRNAs encoding codon-optimized, diproline-modified spike protein from the SARS-CoV-2 Wuhan-Hu-1 strain were produced as previously described^15,16^ with m1Ψ-5-triphosphate (TriLink) instead of UTP and capped cotranscriptionally using the trinucleotide cap1 analog, CleanCap (TriLink), then cellulose purified as previously described.^17^ m1Ψ-containing mRNAs were encapsulated in lipid nanoparticles (LNP) (proprietary to Acuitas Therapeutics) using a self-assembly process as previously described.^18^

### Protein Expression

SARS-CoV-2 Wuhan Hu-1 full length spike (FL-S) protein was purified by Ni-NTA resin from 293F cells transfected with a plasmid that encodes the FL-S (A gift from Florian Krammer, Icahn School of Medicine at Mt. Sinai, New York City, NY).^19^

### ELISA

Immulon 4HBX plates (Thermo Fisher Scientific) were coated with SARS-CoV-2 Wuhan Hu-1 full length spike protein (0.5 μg/mL) diluted in PBS overnight at 40°C. Plates were blocked with 3% bovine serum albumin (BSA; Sigma-Aldrich) in PBS for 1 hour at room temperature and then washed with PBS-T (Phosphate buffered saline [Sigma-Aldrich] with 0.1% Tween-20 [Fisher Bioreagents]). Sera, or monoclonal anti-SARS-Related Coronavirus 2 spike RBD-mFc fusion protein (NR-53796; produced in vitro, BEI Resources, NIAID, NIH), was diluted in 1% BSA in PBS-T and incubated for 2 hours at room temperature. Plates were washed with PBS-T and secondary antibody diluted in 1% BSA in PBS-T was incubated in the plates for 1 hour at room temperature. For IgG quantitation, goat anti-mouse IgG HRP-conjugated secondary antibody (Jackson ImmunoResearch Laboratories, 115-035-071) was diluted 1:1,000 in 1% BSA in PBS-T; for IgM quantitation, goat anti-mouse IgM HRP-conjugated secondary antibody (Jackson ImmunoResearch Laboratories, 115-035-075) was diluted 1:5,000 in BSA in PBS-T. Plates were washed and developed with SureBlue™ TMB 1-component microwell peroxidase substrate (KPL, SeraCare, Milford, MA) for 5 minutes at room temperature and read on a SpectraMAX 190 (Molecular Devices). Data are reported as arbitrary units per milliliter (AU/mL) corresponding to absorbance-matched concentrations to a standard curve of SARS-CoV-2 monoclonal antibody (NR-53796) absorbance. Log transformed values of 0 AU/mL were imputed to half of the threshold of detection (TOD).

### Statistics

Statistical analyses were completed using GraphPad Prism 9. Normal or lognormal distribution of data was confirmed using the Shapiro-Wilk test. Antibody concentrations with skewed distributions were log transformed prior to statistical hypothesis testing. Statistical significance for group comparisons were determined using one-way analysis of variance (ANOVA) with the significance of pairwise comparisons determined by Tukey test (if all possible pairwise comparisons were tested) or Šidák test (if selected pairwise comparisons were tested). All p-values < 0.05 were considered statistically significant. Data are reported as means ± standard deviation (SD) or geometric means with 95% confidence intervals, unless otherwise noted.

## Acknowledgements

We thank all the members of the Hensley laboratory for their thoughtful discussions related to this project. This project has been funded in part with Federal funds from the National Institute of Allergy and Infectious Diseases, National Institutes of Health, Department of Health and Human Services, under Contract No. 75N93021C00015. S.E.H. holds an Investigators in the Pathogenesis of Infectious Disease Awards from the Burroughs Wellcome Fund. R.N.E. was supported by NIH/NIAID T32-AI118684-05 and by the NIH/NIAID Extramural Loan Repayment Program for Pediatric Research (LRP-PR, 1L40AI164510-01).

## Author Contributions

R.N.E, E.M.D, and S.E.H. designed the experiments, analyzed, and interpreted data and wrote the manuscript. R.N.E and E.M.D performed experiments. R.H. and D.W. supervised mRNA-LNP production and provided advice on experimental use of these materials. S.E.H supervised all other activities.

## Data and Materials Availability

The data that support the findings of this study are included in the manuscript. All materials used in this manuscript are available from the authors upon reasonable request.

